# Agonism and grooming behavior explain social status effects on physiology and gene regulation in rhesus macaques

**DOI:** 10.1101/2021.07.16.452731

**Authors:** Noah D. Simons, Vasiliki Michopoulos, Mark Wilson, Luis B. Barreiro, Jenny Tung

## Abstract

Variation in social status predicts molecular, physiological, and life history outcomes across a broad range of species, including our own. Experimental studies indicate that some of these relationships persist even when the physical environment is held constant. Here, we draw on data sets from one such study—experimental manipulation of dominance rank in captive female rhesus macaques—to investigate how social status shapes the lived experience of these animals to alter gene regulation, glucocorticoid physiology, and mitochondrial DNA phenotypes. We focus specifically on dominance rank-associated dimensions of the social environment, including both competitive and affiliative interactions. Our results show that simple summaries of rank-associated behavioral interactions are often better predictors of molecular and physiological outcomes than dominance rank itself. However, while measures of immune function are best explained by agonism rates, glucocorticoid-related phenotypes tend to be more closely linked to affiliative behavior. We conclude that dominance rank serves as a useful summary for investigating social environmental effects on downstream outcomes. Nevertheless, the behavioral interactions that define an individual’s daily experiences reveal the proximate drivers of social status-related differences, and are especially relevant for understanding why individuals who share the same social status sometimes appear physiologically distinct.

## INTRODUCTION

The social environment is an important predictor of health and mortality risk in social mammals (1-3). This observation is particularly well-supported for the relationship between socioeconomic status (SES), physiology, and lifespan in humans, in which low SES predicts higher morbidity and mortality risk across the life course (4, 5). This pattern is to some degree paralleled in studies of dominance rank in other social mammals, in which rank, like SES, reflects systematic, socially structured differences in resource access (6-12). Specifically, in natural mammal populations where a relationship between rank and mortality is observed, higher-ranking individuals tend to live longer lives (3, 13-19), but see (20) for an exception in male baboons). Dominance rank is also a predictor of other aspects of physiology, such that low rank is most commonly (but not always) linked to elevated glucocorticoid levels and increased pro-inflammatory activity (21-26). These patterns again echo a general pattern for SES in humans (27).

Why do low status and high status individuals differ in their physiology and life history? In many cases, access to physical resources and exposure to physical stressors are key contributors. In humans, for example, low SES often predicts poorer health care access and increased food insecurity (28, 29). Smoking, toxin exposure, and exposure to violent crime are also socioeconomically stratified (30-35). The nature of physical stressors differs in other animals, but variation in resource access and physical safety also contributes to status effects on mortality in other species. For example, in cooperatively breeding meerkats, subordinate animals live shorter lives than dominants because they are less spatially central to the group and more likely to leave their groups on extended “excursions.” Increased rate of time away from their groups in turn translates to a higher mortality rate because of increased exposure to predators and reduced foraging success (15). Similarly, in Barbary macaques, dominant individuals are better able to form and/or access larger sleeping huddles, which confer thermoregulatory benefits (36).

However, studies in captivity indicate that social status effects on physiology and health persist even when differences in the physical environment (e.g., diet, housing, predator risk, medical care) are eliminated. In captive long-tailed macaques on an atherogenic diet, low ranking females have been shown to experience a more than two-fold increase in coronary artery atherosclerosis and hyperinsulinemia risk compared to high-ranking females (37-39). In male mice, chronic exposure of subordinate males to dominant males increases the prevalence of atherosclerotic lesions and multi-organ tumors, despite standardized housing and no physical contact between subordinates and dominants (40). Remarkably, these effects translate to a substantial (12.4%) decrease in natural lifespan for subordinate versus dominant male mice (40). Indeed, experimental manipulation of dominance rank itself is sufficient to change glucocorticoid physiology, immune gene regulation, and the response to pathogens in adult female rhesus macaques (25, 41-44). Thus, social status can influence physiological outcomes independent of physical resource limitation, in line with arguments for a contributing role of psychosocial stress exposure to social gradients in health in humans (1, 45).

The basis for these observations must therefore emerge from the quantity, quality, and nature of social interactions themselves. Dominance rank shapes many types of social interactions, providing a potential explanation for why simple measures of hierarchy position have proven so powerful in predicting diverse outcomes (11, 14, 26, 46-50). However, it remains unclear whether dominance rank itself has special explanatory power, or whether it acts as a proxy for a specific subset of social interactions that more directly influence health or fitness outcomes. For example, while dominance rank hierarchies in animals are almost universally estimated using observations of agonistic interactions (i.e., competitive interactions in which a clear “winner” and “loser” can be scored; (11, 51)), individuals that occupy the same hierarchy position can differ in their rate of engagement in agonisms, the proportion of encounters won, or the relative rate of wins versus losses (52). In social primates, where grooming is the primary currency of social affiliation, high-ranking individuals also vary in whether they are more likely to engage in grooming overall (47), more likely to be groomed by subordinates (grooming “up the hierarchy”; (48)), or more likely to target subordinates for grooming (grooming “down the hierarchy”; (53, 54)).

Rank-associated behavioral measures could therefore capture salient variation in the social environment better than dominance rank itself. In support of this possibility, hair cortisol concentrations in a small sample of captive tufted capuchins were positively predicted by rates of aggression received and negatively predicted by grooming network centrality but were not associated with dominance rank (55). In captive rhesus macaques, hair cortisol was also uncorrelated with rank, but was negatively correlated with the rate of initiated affiliative interactions (50). And in wild male chimpanzees, a positive correlation between urinary glucocorticoid levels and high status appears to be primarily explained by rates of aggression, particularly aggression given (56).

Alternatively, dominance rank could serve as a holistic summary of social experience— akin to cumulative adversity or allostatic load indices (57-59) — that captures more information than individual behaviors alone. Distinguishing between these alternatives is important for understanding the mediators of social status effects on health and fitness, including why individuals of the same status often still vary (60). In addition, it may help sharpen our understanding of when dominance rank in nonhuman animals is an appropriate model for socioeconomic status in humans.

To do so here, we leveraged an established nonhuman primate model for studying the causal effects of social status on physiology and gene regulation. We draw on a multi-year study of captive rhesus macaques, in which replicate social groups were formed from adult females with no prior social history of contact. Order of introduction into these groups was the primary predictor of subsequent dominance rank (measured using Elo rating (61)) such that earlier introduction led to higher rank (62). Importantly, because groups were demographically uniform and kept in a controlled diet and housing environment, this paradigm allowed us to focus on the role played by social interactions, in the absence of substantive variation in the physical environment. Both dominance rank and composite measures of behavioral interactions (e.g., overall social engagement (63)) have been shown to predict physiological and gene regulatory outcomes in this study paradigm, but their relative explanatory power has not been compared.

Our analysis drew on multiple data sets, spanning from variation in gene regulation to differences in cortisol levels and mitochondrial DNA copy number (25, 41-43, 64). Using these data, we addressed three questions. First, we asked how the explanatory power of simple and composite measures of affiliative and agonistic social interactions compares to that of dominance rank. Second, we asked whether the phenotypes for which rank-associated agonistic interactions have the most predictive power differ from those in which rank-associated affiliative interactions are most important. Third, we asked whether variation in social interactions accounted for heterogeneity in the effects of dominance rank itself. This analysis allowed us to investigate whether individual-specific social experience explains why individuals differ in response to the same position in a status hierarchy.

## METHODS

### Study subjects

Study subjects were 45 adult female rhesus macaques housed in nine social groups (n = 5 per group) at Yerkes National Primate Research Center (Figure 1A; note that the sample size of unique individuals ranges from 42-45 across data sets: Table S1). To initially form each group, females were sequentially introduced into 25 m x 25 m indoor-outdoor run housing over the course of 2 – 15 weeks in January – June 2013. Following introduction of all group members, we monitored dominance rank and behavioral interactions in each group using focal sampling (223.5 hours of focal observation data collected across ∼1 year of observations; (42)). We refer to data and samples collected during this time period as study “Phase 1.” Social group membership was then rearranged for study “Phase 2,” beginning in March – June 2014. In this rearrangement, study subjects were sequentially introduced into new groups, where group membership was composed of females who had occupied the same or adjacent ordinal ranks in Phase 1. This approach allowed us to observe the same females after most switched positions in the rank hierarchy between Phase 1 versus Phase 2: half became higher-ranking in Phase 2, while the other half became lower-ranking in Phase 2. We collected an additional 121.5 hours of focal sampling data over a ∼11 month time period following the rank rearrangement. Throughout both phases of the experiment, all study subjects were maintained on unrestricted access to a typical low-fat, high-fiber nonhuman primate diet (Purina Mills International, LabDiets, St. Louis, MO).

**Figure 1.**
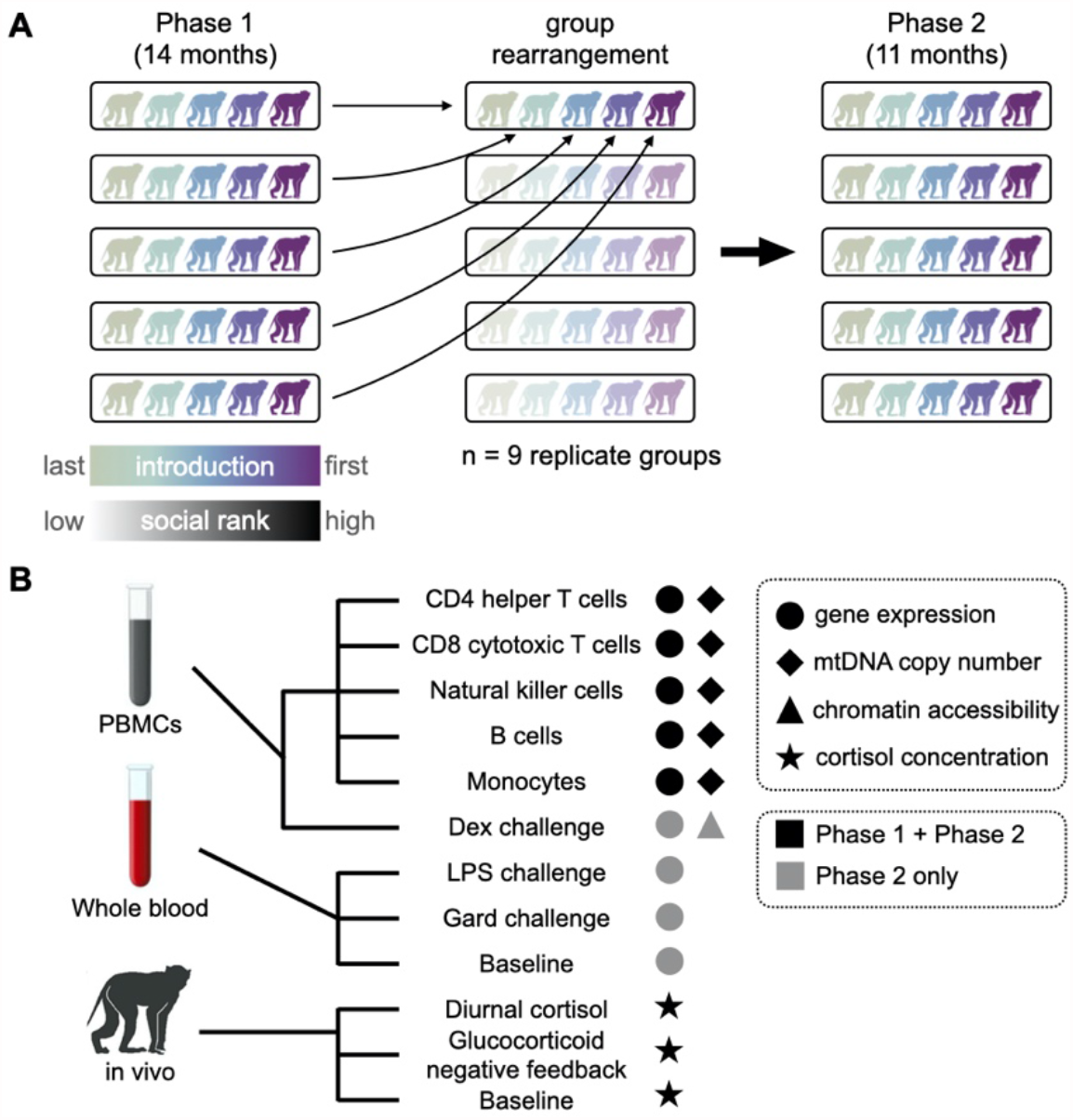
Study design and data sets. (A) Schematic illustration of experimental design. In both Phase 1 and Phase 2 (post group rearrangement), the order of introduction is a strong predictor of social rank in the dominance hierarchy. B) Schematic overview of data sets re-analyzed in this study.

### Data sets

We re-analyzed data on five sets of outcome variables, from five published studies (Figure 1; (25, 41-43, 64)). For each data set we followed the low-level quality control and processing steps described in the original published paper as closely as possible, with some modifications to accommodate the goals of this study (see Supplementary Methods and Table S2 for sample metadata). Data sets included:

i. gene expression (RNA-seq) data from five flow cytometry-sorted cell types purified from peripheral blood mononuclear cells: CD4^+^ helper T cells, CD8^+^ cytotoxic T cells, natural killer (NK) cells, B cells, and monocytes. Gene expression data were collected for each individual, for each cell type, in both Phase 1 and Phase 2 (see (42) for information on flow cytometry and cell sorting strategy).
ii. gene expression (RNA-seq) data from white blood cells purified from a whole blood *in vitro* challenge experiment. Aliquots of cells from each study subject were collected after exposure to cell culture media only (non-stimulated), 1 μg/mL ultra-pure lipopolysaccharide (LPS) from the *E. coli* 0111:B4 strain (LPS), or 1 μg/mL Gardiquimod (Gard), in Phase 2 only.
iii. gene expression (RNA-seq) and chromatin accessibility (ATAC-seq) data from peripheral blood mononuclear cells (PBMCs) purified from a PBMC *in vitro* challenge experiment. Aliquots of cells from each study subject were collected after exposure to cell culture media and either 0.02% EtOH (Dex vehicle) or 1 μM dexamethasone (Dex), in Phase 2 only.
iv. serum cortisol concentration, measured in Phase 1 and Phase 2 to capture: (a) diurnal endogenous cortisol levels across the day, at 0800, 1100, and 1700 hours; (b) diurnal cortisol slope, based on the same three measurements; and (c) glucocorticoid negative feedback, based on endogenous cortisol levels at 1.5, 4.5, and 24 hours following *in vivo* injection with the synthetic glucocorticoid Dexamethasone;
v. mitochondrial DNA (mtDNA) copy number from Phase 1 and Phase 2, for each of the five flow cytometry-sorted cell types in (i). mtDNA copy number was assessed using qPCR against the rhesus macaque mitochondrial genome and normalized against a single-copy locus in the nuclear genome.

### Behavioral data and hierarchy characteristics

All behavioral data were derived from focal observations conducted in 30-minute blocks (223.5 hours in Phase I and 121.5 hours in Phase II; (42)). Because of the small size of our study groups and due to the layout of group run housing, we were able to conduct group-level focal observations throughout the study. During each focal observation, trained observers collected data on agonistic, affiliative, self-directed, and sexual behaviors following an established ethogram (65). We classified threats, attacks, and chases as agonistic interactions (where withdrawals and grimaces indicated the loser of an agonistic interaction), and further distinguished between agonistic interactions that involved physical contact (e.g., slaps, bites) and the much more common interactions that reinforce status but occur without physical contact. Non-contact agonistic interactions accounted for ∼94% of all agonistic interactions.

We compared the predictive power of dominance rank to simple summaries of the rates of social interactions and to composite measures based on multiple behavioral measures. To estimate rank, we used the Elo rating, a continuous measure that updates following every scorable interaction and where a higher Elo rating corresponds to a higher rank in the dominance hierarchy (61, 66). Elo ratings were calculated in the R package *EloRating* (66), using default parameters for the starting Elo value (1000) and k (100), the parameter that determines the maximum change in rating following an interaction. As simple summaries of competitive social interactions, we calculated the overall rate, in events per hour, that a female was the target of an agonistic interaction (“agonism received”) and the overall rate that a female targeted other group mates (“agonism given”). We also calculated agonism rates for noncontact and contact interactions separately, for a total of six measures of simple agonism rates. To measure affiliative interactions, we calculated rates of grooming received and grooming given, in minutes per hour. Finally, we considered three composite measures of social interactions: (i) the difference in the rate of overall agonisms given versus received (“agonism asymmetry;” large positive values tend to be characteristic of higher ranking animals); (ii) the first principal component (“PC1.ag”) estimated from all six simple measures of agonism rates, plus agonism asymmetry; and (iii) the first principal component of the grooming rate data, considering both grooming given and grooming received (“PC1.groom”).

All behavioral variables were correlated with Elo, although the strength of that correlation varies across behavioral measures and data sets (from r = 0.38 for grooming given rate in Phase 1, to r = 0.86 for agonism asymmetry in Phase 2; Figure S1). Overall, higher-ranking animals tended to both give and receive more grooming, received less aggression from other group mates, and engaged in agonistic behavior targeted towards other group mates at a higher rate (Figure S1). Agonism-related measures were more closely correlated with Elo than grooming measures, as expected since Elo is based only on the outcomes of agonistic, not affiliative, interactions.

Because estimation of dominance rank depends only on *consistency* in the directional outcomes of agonistic interactions, dominance rank estimates may be less noisy and/or less temporally heterogeneous than measures of behavioral rates. This difference would arise, for example, in hierarchies in which agonism rates are low except during brief periods of instability or hierarchy challenges. Female rhesus macaque hierarchies are highly stable over time, so high variance is unlikely to compromise our measures of behavioral rates. However, before modeling the behavioral rate data, we assessed the stability of our behavioral rate data. To do so, we split the data in half by alternating observation days, so that each half was equal in size and represented the entire collection range. We then calculated Pearson’s correlation coefficient between the two data splits, for each behavioral variable, where the expectation is that consistent behavioral rates would produce high correlations between the data splits. Finally, we assessed the steepness (67), implemented in the R package *steepness* (v0.2-2; (68)), linearity (h’; (69)), directional consistency (the proportion of contests won by higher-ranking individuals against lower-ranking individuals), and stability (S; implemented in R package *EloRating* (66)) of each group hierarchy. All hierarchy statistics were calculated using R package *compete* (v0.1, (70)), unless otherwise noted.

### Modeling the effects of rank and social interactions

To compare the effects of Elo rating and social interaction measures on each outcome variable, we replicated the modeling approach described in the original analyses of each data set as closely as possible (Supplementary Information). For the cortisol data sets, we modified the original models because they had originally been built to test the effects of behavioral tendencies (dimensions of “personality”), which were derived from the social interaction data that we wished to compare with Elo rating (25). To avoid collinearity and increase comparability to the other data sets, we therefore removed measures of behavioral tendencies from the cortisol level models. For all data sets, we modeled the outcome variables of interest using linear models or linear mixed effects models, controlling for relatedness, age, batch, and other technical covariates specific to each data set. For the genomic data sets, we evaluated each gene or chromatin accessibility window separately.

For each outcome variable, we fit both an “Elo” model that tested for an association between rank and the outcome, as well as a series of alternative models that substituted one of the measures of social interactions (six measures of agonism rates, two measures of affiliation rates, and three composite measures) for Elo, but otherwise retained an identical model structure. For the gene expression and chromatin accessibility data sets, we controlled for multiple hypothesis testing using a false discovery rate approach and FDR threshold of 0.1, calibrated against permutation-derived empirical null distributions (100 permutations per data set).

For a subset of the flow-sorted gene expression data sets, we also asked whether the residual variance in gene expression that was unexplained by rank could be explained by agonism asymmetry. This model tested for an association between residual agonism asymmetry, adjusted for Elo rating, and residual gene expression, also adjusted for Elo rating. As with the previous gene expression analyses, we controlled for multiple hypothesis testing using an FDR threshold of 0.1, calibrated against a permutation-based null (100 permutations per data set). Finally, we tested whether enrichment of Molecular Signature Database (MSigDB) Hallmark gene sets (71) differed for rank-associated genes versus behavioral rate-associated genes in the flow-sorted gene expression data sets using the R package *fgsea* (72), with default parameters.

## RESULTS

### Hierarchy characteristics and social interaction summaries

As previously reported for the rank manipulation experiment (42), dominance ranks based on Elo ratings were strongly predicted by the order that females were introduced into their groups (Phase 1: r = −0.57, p = 4.1×10^−5^; Phase 2: r = −0.68, p = 3.3×10^−7^). Grooming and agonism rates also tracked changes in Elo rating between phases (Figure 2). Once formed, hierarchies were highly linear (mean h’ = 0.99 ± 0.01 s.d.; n = 18 groups across both study phases) and the outcomes of agonistic encounters were highly directionally consistent (median proportion of contests won by higher-ranking individuals = 0.95 ± 0.09 s.d.). Group dominance hierarchies were also very stable within each phase (median S was the same for both phases: 0.997 ± 0.002 s.d.; a value of 1 means that rank changes never occur). We observed somewhat greater variation in hierarchy steepness (median: 0.88 ± 0.08 s.d.), such that high-ranking and low-ranking animals in some groups were more strongly differentiated than others, but this reflected a quantitative, not qualitative, distinction. We observed no significant differences in group hierarchy characteristics (linearity, directional consistency, stability, and steepness) between the two phases (all p > 0.05).

**Figure 2.**
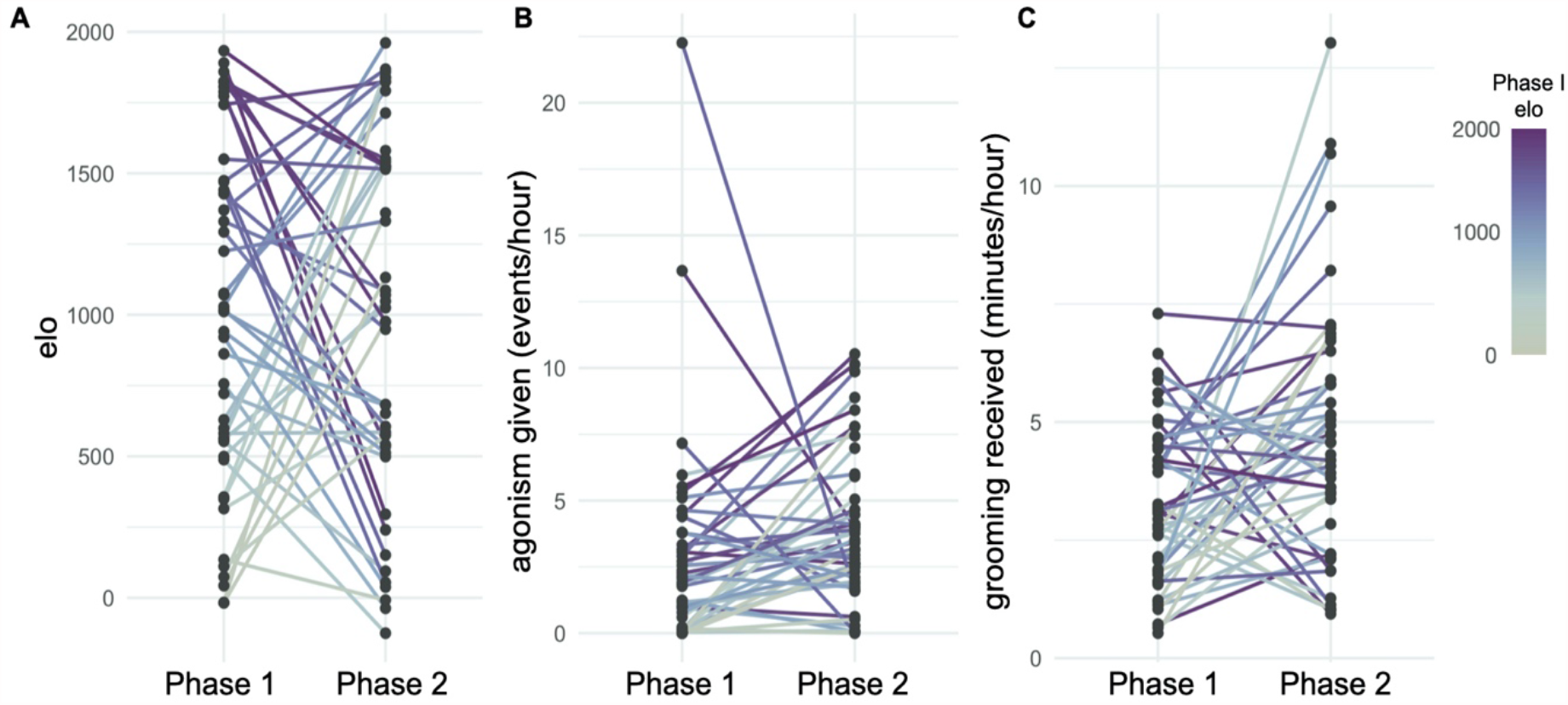
Rank and behavioral patterns change across study phases. **(A)** Dominance rank (Elo) shifts after the mid-study rank rearrangement; approximately half of the study subjects moved up the hierarchy and half moved down. **(B)** Raw agonism and **(C)** grooming rates also shift between study phases, following changes in rank (agonism given and grooming received are shown here, but patterns are similar for agonism received and grooming given). Lines connecting dots between Phase 1 and Phase 2 show change for the same female. Correlations between values in Phase 1 and Phase 2 are all p > 0.05.

Together, these summaries are consistent with the clearly delineated, “despotic” hierarchies typical of rhesus macaque females (73), despite the fact that the process of hierarchy establishment in this experimental setting differs from the typical process of matrilineal inheritance in unmanipulated groups. Consistent with the data on dominance rank, our behavioral measures were also consistent across data subsamples: correlations between split data sets were highly significant across all behavioral measures (all p < 2.5 × 10^−14^; Figure S2).

### Agonism rates often predict gene regulatory phenotypes better than dominance rank

We considered nine genome-wide gene expression data sets and one genome-wide chromatin accessibility data set. For eight out of the nine gene expression data sets (all but NK cells), a simple or composite measure of agonism rates identified more significantly associated genes than identified using parallel models based on dominance rank (FDR < 0.1) (Figure 3A; Tables S3-S5). In contrast, the predictive power of grooming-related behaviors was consistently lower than for Elo (Figure 3A). For data sets in which dominance rank affects a large number of genes, such as LPS- and Gard-challenged white blood cells, the difference between rank and agonism rates was minor. For example, dominance rank drove differential gene expression for 6,234 genes (FDR < 0.10) in LPS-stimulated white blood cells, but agonism asymmetry was associated with differential expression for n=6,595 genes (5.79% more).

**Figure 3.**
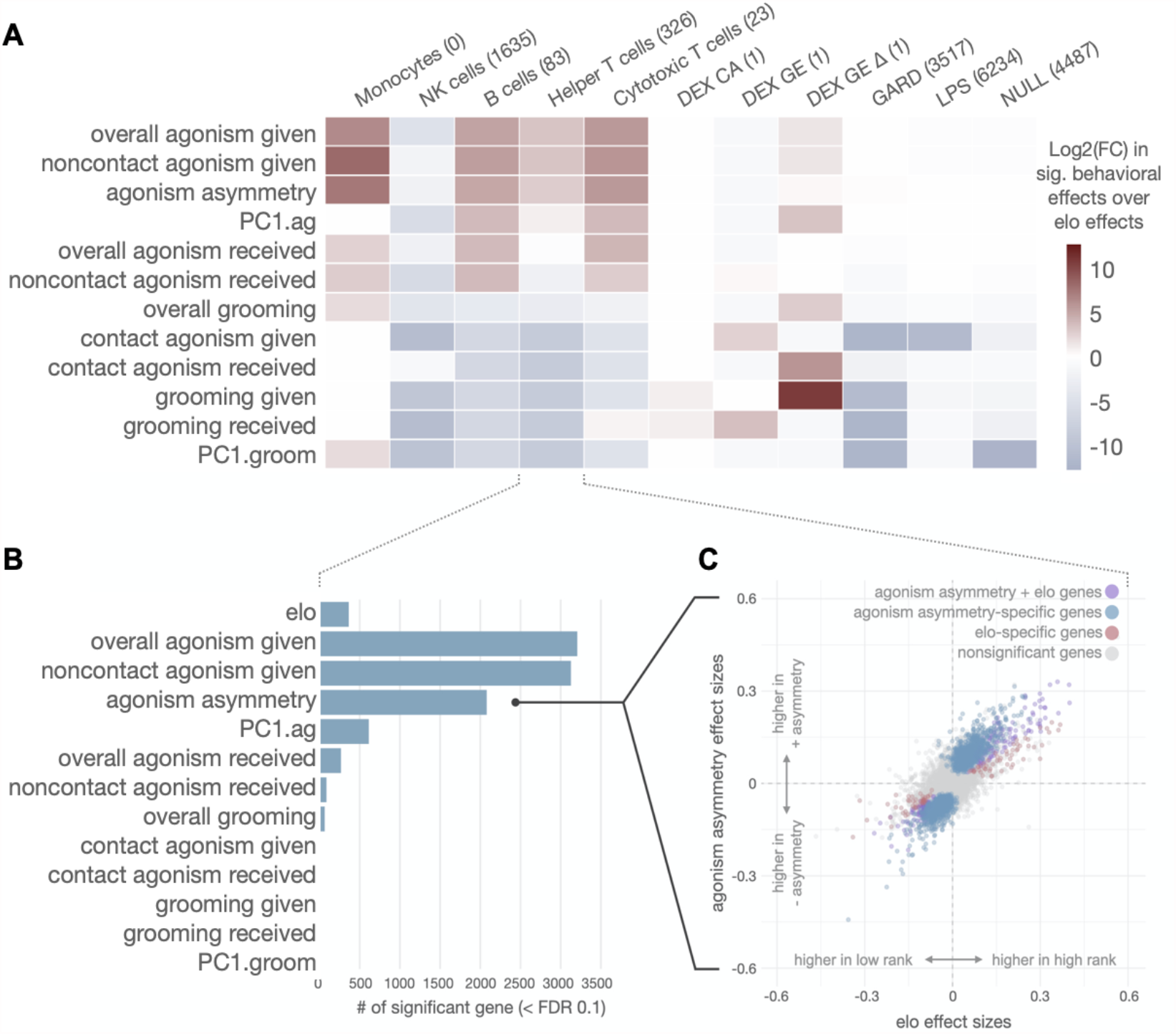
Behavioral summaries often explain gene expression patterns better than dominance rank. **(A)** Foldchange difference in the number of significant predictor variable-gene expression associations, relative to those identified using Elo (red = more associations; blue = fewer associations). Each data set is represented in a separate column, with the number of Elo-associated genes for that data set in parentheses above the data set label. **(B)** As an example, in CD4^+^ helper T cells, four agonism-related behavioral variables identify more social environment-associated genes than dominance rank (Elo). **(C)** Overall effect size estimates are highly correlated between Elo ratings and agonism summaries (e.g. Elo-agonism asymmetry correlation *R*^*2*^ = 0.68, p < 2.7e-14; see also Figure S3); however, while relatively few genes show Elo rating-specific effects (red dots), agonism asymmetry identifies many genes that are not identified by analyzing Elo (blue).

The difference was more striking for gene expression data sets in which few significant rank-associated genes were detected (Figure 3A; Tables S3-S5). For example, in purified B cells, dominance rank explained variation in the expression levels of 83 genes, but agonism asymmetry identified significant associations with 2,498 genes. Similarly, in purified CD4^+^ helper T cells, dominance rank explained variation in the expression levels of 326 genes, but agonism asymmetry identified significant associations with 2,088 genes, a 6.4-fold increase (Figure 3B-C). Finally, in monocytes, modeling Elo rating identified no effect of dominance rank, but modeling agonism asymmetry identified 152 genes, highlighting an effect of the social environment that would be missed by considering dominance rank alone. Notably, though, in nearly all cases in which both Elo and agonism rate effects were detected, the directionality of these effects was consistent (i.e., if a gene was more highly expressed in high-ranking animals, it was also more highly expressed in animals who directed more agonisms to others than they received, and vice-versa). This observation, as well as the generally high correlation between Elo effect sizes and agonism asymmetry effect sizes (Figure 3C; Figure S3) indicate that the increased predictive power for agonism rates reflects a quantitative, not a qualitative, difference.

Overall, we observed the greatest power increase, relative to Elo, when modeling agonism asymmetry or simple measures of overall or non-contact agonisms given (because most agonistic interactions are non-contact, these measures are highly correlated: r = 0.88; p = 2.2 × 10^−16^). To a lesser degree, we also observe a power increase for an alternative composite measure, the first principal component of all agonism rate measures (PC1.ag), and for rates of agonisms and noncontact agonisms received. In contrast, measures of contact aggression rates (either given or received) are poor predictors of gene expression variation. This result may arise because contact aggression is rarer and more episodic, or because frequent use of contact aggression tends to be idiosyncratic to specific individuals (e.g., rates of contact aggression given among alpha females range from 0 – 0.9 events per hour). Low and variable expression of contact aggression may reduce its biological significance to changes in gene regulation, as well as the accuracy of our rate estimates.

In contrast to the gene expression data sets measured in purified PBMC subtypes or in immune challenge experiments, we observed little to no improved power for agonism rates in gene expression and chromatin accessibility data measured in the glucocorticoid challenge experiment. However, for the fold-change gene expression *response* to Dex (i.e., difference between Dex-challenged and control cells from the same individual), rates of grooming given were associated with 2,895 genes at a 10% FDR, compared to only 1 at the same statistical threshold for Elo rating. We observed no strong explanatory power for either dominance rank or social interaction measures when analyzing chromatin accessibility patterns for the same data set.

### Affiliative social interactions often predict glucocorticoid and mitochondrial DNA phenotypes better than dominance rank

Our results for the gene expression data sets indicate that measures of social interactions often predict downstream outcomes better than rank itself. To test this possibility further, we re-analyzed a data set from the same animals on diurnal cortisol levels, cortisol slope, and cortisol concentration 1.5, 4.5, and 24 hours post-dexamethasone challenge (Dex induces a negative feedback loop that reduces the production of endogenous cortisol). We also re-analyzed data on mitochondrial DNA copy number, which has been linked to social interaction, stress, and depression in rhesus macaques, mice, and humans (74-77).

Three measures of grooming behavior (overall grooming rates, grooming given, and the first principal component of all grooming measures), but no measures of agonism rates, significantly predict diurnal cortisol slope (Table 1; Table S6). Specifically, females who engaged in less grooming behavior had blunted diurnal changes in cortisol (smaller slopes) than those who engaged in more grooming behavior. Models with each of these three variables also performed better than an alternative model where Elo rating was substituted for grooming rates and better than the null model (|ΔAIC| > 2). We observed no other significant effects of dominance rank, affiliative behavior, or agonistic behavior on diurnal cortisol or cortisol levels after Dexamethasone challenge, with the exception of contact aggression given at the 24-hour time point. However, while contact aggression was nominally significant (p = 0.047), the model including contact aggression did not perform better than the Elo model (ΔAIC = −0.86). Finally, the rate of grooming received and a composite summary of grooming (PC1.groom), but not agonism rates or dominance rank, were significant predictors of mtDNA copy number (t = 2.36, p = 0.019 and t = −2.43, p = 0.015, respectively; Table S7). While neither behavioral variable performed better than the null model, both grooming received and PC1.groom did improve on the Elo model (Δ-AIC = - 5.50 and Δ-AIC = −5.61, respectively).

**Table 1.** Comparison between the effects of rank and significant behavioral variables for cortisol and mtDNA copy number data sets.

| **Behavioral Variable<sup>1</sup>** | **Outcome** | **beta** | **SE** | **t** | **p** | **AIC** | **Δ-null** | **Δ-Elo** |
| --- | --- | --- | --- | --- | --- | --- | --- | --- |
| Null Model | cortisol slope |  |  |  |  | 46.68 |  | -1.02 |
| Elo |  | 0.066 | 0.070 | 0.949 | 0.347 | 47.70 | 1.02 |  |
| ***PC1.groom*** |  | 0.106 | 0.051 | 2.082 | 0.044 | 44.15 | -2.53 | -3.55 |
| ***grooming given*** |  | 0.133 | 0.062 | 2.162 | 0.036 | 43.82 | -2.86 | -3.88 |
| ***overall grooming*** |  | 0.148 | 0.068 | 2.197 | 0.034 | 43.67 | -3.01 | -4.03 |
| Null Model | 24 h post-Dex |  |  |  |  | 219.27 |  | -1.63 |
| Elo |  | 0.864 | 0.471 | 1.833 | 0.074 | 217.64 | -1.63 |  |
| *contact agonism given* |  | 1.688 | 0.824 | 2.048 | 0.047 | 216.78 | -2.49 | -0.86 |
| Null model | mtDNA copy number |  |  |  |  | 395.97 |  | -6.7 |
| Elo |  | -0.003 | 0.038 | -0.09 | 0.93 | 402.67 | 6.7 |  |
| ***PC1.groom*** |  | -0.083 | 0.034 | -2.43 | 0.015 | 397.06 | 1.09 | -5.61 |
| ***grooming received*** |  | 0.09 | 0.038 | 2.36 | 0.019 | 397.17 | 1.20 | -5.50 |

<sup>1</sup>Italicized variables are nominally significant at $p < 0.05$ ; italicized and boldfaced variables are nominally significant and improve on both the null model and Elo models ( $|\Delta AIC| > 2$ ).

### Relationship between rank and behavioral effects on gene expression

To investigate whether the behavioral data capture information about social environmental effects on gene expression, beyond that captured by dominance rank, we focused on our agonism asymmetry measure and data from three of the five flow-sorted PBMC subpopulations (excluding monocytes, where we did not detect any significant rank-gene expression associations, and NK cells, where we identified more Elo-associated genes than agonism asymmetry-associated genes). In all three data sets, Elo-associated genes were largely nested within agonism asymmetry-associated genes (68.7% [helper T cells] – 96.4% [B cells] of Elo-associated genes were also detected as agonism asymmetry-associated, Figure S4). The set of Elo-associated genes and agonism asymmetry-associated genes thus significantly overlap (helper T cells log2(OR): 2.06; cytotoxic T cells log2(OR): 4.19; B cells log2(OR): 4.26; all p < 2.0 × 10^−16^). However, the stronger signal of agonism asymmetry reveals several enriched pathway/gene sets that are not detectable in the rank analysis. Specifically, in helper T cells and cytotoxic T cells, we identify significant (false discovery rate < 0.1) enrichment for 10 Hallmark gene sets that are not detected for rank-associated genes (Table S8).

Agonism asymmetry-specific enrichments in CD4 T cells include two key innate immune signaling gene sets (the interferon gamma response and interferon alpha response: [ES = −0.34, FDR = 0.002 and ES = −0.39, FDR = 0.003 respectively, versus ES = −0.21, FDR = 0.56 and ES = −0.20, FDR = 0.79 for Elo]). All three gene sets are enriched for higher expression in individuals with high agonism asymmetry.

Finally, for rank-associated genes in helper T cells and B cells, we found that agonism asymmetry explained residual variance unexplained by rank. Specifically, levels of agonism asymmetry (relative to the expected value based on rank) significantly predicted rank-adjusted gene expression levels for 18.1% (59 genes < FDR = 0.1) and 28.9% (24 genes < FDR = 0.1) of rank-associated genes in helper T cells and B cells, respectively (Table S9). Thus, variation in gene expression levels for individuals of the same rank is often partially explained by variation in the behavioral interactions those animals experience day-to-day.

## DISCUSSION

Together, our findings indicate that simple measures of agonistic and affiliative social interactions in female rhesus macaques often have greater predictive power for gene expression and glucocorticoid phenotypes than Elo ratings, even though these behavioral measures are moderately to strongly correlated with Elo. Our results are consistent with previous work using a similar paradigm, in which reduced social engagement predicts compromised glucocorticoid negative feedback (63). Indeed, for rank-associated gene expression traits, engaging in a higher (or lower) rate of agonistic interactions than expected based on rank accounts for additional variance unexplained by rank itself—highlighting one possible explanation for heterogeneity in the predictive power of dominance rank across individuals, groups, and species (21-26). Our results therefore suggest that analyzing behavioral correlates of rank offers insight into the social determinants of phenotypic variance beyond that obtainable from analyzing dominance rank alone.

These findings emphasize that, while dominance rank structures social interactions and resource access, rank itself is not the proximate driver of phenotypes that follow a social gradient. Instead, associations between rank and downstream outcomes capture differences in the physical and social resources that flow to animals who occupy high (versus low) rank. As our findings emphasize, these resources can take many forms, including control of one’s social environment, and measuring dominance rank alone does not necessarily reveal the nature of those resources. Indeed, in humans, sociologists have long argued that socioeconomic status is a “fundamental cause” of differences in health, irrespective of disease outcome or intervening mechanism (29). This occurs because high SES confers better access to whatever resources are advantageous in a particular environment or context (e.g., better nutrition in some environments, reduced exposure to infectious disease in others). Dominance rank has parallel implications in nonhuman animals, although the benefits to rank are better viewed through the lens of Darwinian fitness instead of health, especially given costs to high status in some species-sex combinations (20, 56, 78-80).

Despite the improved power of behavioral summaries in our analyses, rank remains a valuable summary of social experience for several reasons. First, the generality of rank as a measure of competitive advantage can be a feature for some types of analysis (e.g., in comparative work, where the specific benefits of high status differ across species, populations, or sexes). Second, dominance rank may be less vulnerable to measurement error. Quantifying rank requires estimating the directional outcome of competitive interactions: that is, which animal will win an encounter, conditional on its occurrence. Measures of behavioral rates, on the other hand, require accurately estimating the frequency with which a behavior occurs, which can be challenging for episodic behaviors, in low visibility environments, or when behavioral observations are sparse. Third, directly testing intervening behavioral pathways is challenging when strong candidate pathways are unclear. In natural populations, especially, dominance rank structures access to both physical and social resources, complicating direct assessment of mechanism without a strong *a priori* hypothesis.

Behavioral measures can nevertheless help refine our understanding of social status effects. For example, in Kibale National Park, Uganda, high ranking chimpanzee males exhibit higher urinary glucocorticoid levels than low ranking males (56). This relationship appears to be explained by rates of aggression directed towards other males, which, in this population, is more variable across the dominance hierarchy than rates of aggression received. The authors thus propose that energetically expensive, protracted displays mediate the relationship between rank and glucocorticoid physiology in male chimpanzees, highlighting a cost to dominance (56). Behavioral data may also highlight why the effects of rank on physiological measures vary across social groups and populations. A number of studies, for instance, suggest that rank effects are exacerbated (and sometimes directionally reversed) during periods of rank instability (e.g., (81-83). Elevated rates of aggression are likely to be a proximate explanatory factor.

Finally, our results point to an intriguing pattern in which the molecular and physiological imprint of rank-associated affiliative behavior may differ from that of rank-associated competitive interactions. In our study system, both grooming and agonisms are structured by dominance rank, making them difficult to fully disentangle. However, our results dovetail with recent findings in wild baboon females, which indicate distinct signatures of dominance rank versus social bond strength on immune cell gene expression (84). Assuming that high status and affiliative behavior are both beneficial, high status animals may therefore be doubly advantaged in settings where high rank is a primary predictor of social integration. In settings where social affiliation and social bond formation are partially independent of rank, however, affiliative behavior may provide opportunities for social buffering against status-related costs (85). Notably, while analyses that take into account both aspects of the social environment are now common in studies of glucocorticoid physiology and, to a lesser extent, natural lifespan, studies of molecular variation (e.g., gene expression, DNA methylation) have generally focused only on one (41, 42, 86, 87). Our findings here suggest that using behavioral measures to dissect the social environment into its component parts can further refine these types of analysis.

## DATA ACCESSIBILITY

All data analyzed here were previously published under the following accession numbers: flow-sorted gene expression data and LPS challenge data are available under GEO accession number GSE83307; Dex challenge data are available under NCBI BioProject ID PRJNA476378; Gard challenge data are available under GEO accession number GSE136124; glucocorticoid data are available in the supplemental materials of (25); mtDNA data are available in the supplemental materials of (64). Code and processed behavioral data used in this study can be found at https://github.com/ndsimons/MacaqueBehaviorEffects2021.

## ACKNOWLEDGEMENTS

We thank J. Whitley, A. Tripp, N. Brutto, and J. Johnson for maintaining the study subjects and collecting behavioral data. We thank N. Snyder-Mackler, J. Kohn, and R. Debray for assistance with analysis of previously generated data sets, C.R. Campbell for assistance with code review, and P. Maurizio, J. Batista, and other members of the L.B.B. and J.T. laboratories for helpful discussions. This work was supported by NIH grants F32AG062120, R01GM102562, and R01AG057235, and by high-performance computing resources from the North Carolina Biotechnology Center (Grant Number 2016-IDG-1013).

