## Supplemental Materials for "Agonism and grooming behavior explain social status effects on physiology and gene regulation in rhesus macaques"

**Table of Contents**

1. **Supplementary Methods**
   a. Data Provenance & Statistical modeling
      i. Flow-sorted gene expression data sets
      ii. Lipopolysaccharide and Gardiquimod challenge gene expression data sets
      iii. Dexamethasone challenge gene expression and chromatin accessibility data sets
      iv. Cortisol-related data sets
      v. mtDNA copy number data set
2. **Supplementary Tables (Tables S2 – S8 are large and contained as tabs in a separate Excel file)**
   a. Table S1: Data sources and sample sizes
   b. Table S2: Sample metadata
   c. Table S3A-E: Flow-sorted PBMC gene expression model results
   d. Table S4A-C: LPS- & GARD-challenged gene expression model results
   e. Table S5A-C: DEX-challenged gene expression and chromatin accessibility model results
   f. Table S6: Glucocorticoid level model results
   g. Table S7: mtDNA copy number model results
   h. Table S8A-D: Gene set enrichment results
   i. Table S9: Residual agonism asymmetry model results
3. **Supplementary Figures**
   a. Figure S1. Relationships between Elo and behavioral variables
   b. Figure S2. Behavioral rate estimates are consistent across subsampled data sets
   c. Figure S3. Effect size relationships between Elo and other behavioral variables.
   d. Figure S4. Overlap between Elo-associated and agonism asymmetry-associated genes

**SUPPLEMENTARY METHODS**

**Data Provenance & Statistical Modeling**

For all data sets, we followed the modeling approach reported in the original study, except where modifications were necessary to accommodate the goals of this study. The statistical methods are described below in brief, but are also outlined in additional detail in the original papers (Table S1).

*Flow-sorted gene expression data sets*

We analyzed five data sets on cell type-specific gene expression from (1) (available at https://github.com/nsmackler/status_genome_2016 and GEO accession number GSE83307). Each data set includes samples derived from purified peripheral blood mononuclear cells and physically sorted into individual PBMC components on a BD FACSAria IIu, followed by genome-wide gene expression quantification using RNA-seq. Samples were collected in both Phase 1 and Phase 2 of the study. We analyzed five PBMC subsets for each sample, which corresponded to classical monocytes (CD3^-^/CD20^-^/HLA-DR^+^/CD14^+^), natural killer cells (CD3^-^/CD20^-^/HLA-DR^-^/CD16^+^), B cells (CD3^-^/CD20^+^/HLA-DR^+^), helper T cells (CD3^+^/CD8^-^/CD4^+^), and cytotoxic T cells (CD3^+^/CD8^+^/CD4^-^). This study design resulted in a total sample size of 86 - 89 for each of the five cell types (maximum of 2 samples per cell type per study subject). We followed the sample inclusion/exclusion rules reported in the original paper.

We analyzed data from each cell type separately but considered data from both phases together, as in (1). Gene expression data were represented as batch-corrected residuals of *voom*-normalized gene counts (2). For each cell type, we modeled each normalized gene expression phenotype separately, as follows, considering 8,388 – 8,951 gene expression phenotypes per cell type:

$$y = \mu+b\beta_{b}+a\beta_{a}+g+\varepsilon,$$

$$g \sim MVN(0,\sigma_{u}^{2}K),$$

$$\varepsilon\sim MVN(0,\sigma_{\varepsilon}^{2}I)$$

where $y$is an $n x 1$vector of normalized gene expression levels for $n$ samples across both phases; $\mu$ is the intercept; $b$ is an $n$ x 1 vector for a given behavioral variable (e.g., Elo, agonism asymmetry, grooming given) and $\beta_{b}$ is its effect size; $a$ is an $n$ x 1 vector of age at the time of sampling and $\beta_{a}$is its effect size; $g$ is an *n* x $1$ vector for a multivariate normally distributed (*MVN*) random effect that captures genetic non-independence (e.g., due to relatedness) among samples and individuals; and $\varepsilon$ is an $n$ x 1 vector of residual error. *σ_g_^2^* and *σ_e_^2^* are the genetic and environmental variance components, respectively, and the relatedness matrix, *K*, is constructed from genotypes called using GATK’s Best Practices for genotyping RNA-seq data (<https://gatk.broadinstitute.org/hc/en-us/articles/360035531192?id=3891>) (3). $I$ denotes the identity matrix.

*Lipopolysaccharide and Gardiquimod challenge gene expression data sets*

We also analyzed RNA-seq gene expression profiles from paired baseline control (NULL: n = 40), LPS-challenged (LPS: n = 43), and Gardiquimod-challenged (GARD: n = 42) samples collected in Phase 2 of the study, as reported in Snyder-Mackler et al. 2019 (4) and Sanz et al. 2020 (5) (data available under NCBI BioProject ID PRJNA476378 and GEO accession number GSE136124, respectively). We analyzed the LPS and control data together using mixed effects models similar to those used for the flow-separated cell types (see above) but included a fixed effect of condition and nested the effects of rank or the behavioral variable of interest within condition (control or LPS). We also nested the effects of age at the time of sampling and the first two principal components of cell type composition within condition. Estimates for both the LPS data set and the control (null) data set are based on this nested model, following (1).

GARD estimates are based on RNA-seq data from biological samples that were collected during the same draw as the control and LPS-challenged samples Sanz et al. 2020 (5). We therefore analyzed the effects of Elo and each behavioral variable of interest in the GARD data set using the same model structure as for the control and LPS-challenged data. However, to avoid generating two estimates of these effects in the control (NULL) condition, we analyzed gene expression data from the GARD challenge condition in a model excluding the control samples (i.e., a non-nested model). We therefore removed the fixed effect of condition (as there was no variation in challenge condition in these models), but retained controls for age, the first two principal components of tissue composition, and genetic structure.

*Dexamethasone challenge gene expression and chromatin accessibility data sets*

We analyzed dexamethasone (DEX)-challenged PBMC gene expression and chromatin accessibility data sets from (4) (available at https://github.com/nsmackler/Dex_status_2018 and GEO accession number GSE83307). Each data set included n = 43 individuals from Phase 2 of the experiment. We modeled the effects of Elo and each behavioral variable on three DEX-related outcome variables: gene expression following DEX challenge (DEX GE), chromatin accessibility following DEX challenge (DEX CA), and the gene expression *response* to DEX, measured as log_2_ fold-change difference between the baseline and DEX-challenged state ($\Delta$DEX GE). For each gene (for gene expression levels) and genomic window (for chromatin accessibility), we modeled normalized, batch-corrected expression/ATAC-seq count data as the response variable in a mixed effects model with a structure that paralleled the other gene expression analyses. Following the original study, we controlled for genetic relatedness and cell type composition using the first three principal components from flow cytometry-based phenotyping data (4).

*Cortisol-related data sets*

We analyzed diurnal cortisol concentration data from (6) (available in the original paper Supplementary Information). Here, serum from 45 individuals was used to measure serum cortisol concentrations, either based on endogenous levels throughout the day (at 0800 and 1700 hours) or after administration of Dexamethasone, a synthetic glucocorticoid that leads to suppression of endogenous cortisol production (DEX suppression test, which assays glucocorticoid negative feedback). We estimated the effects of Elo or each behavioral variable independently on endogenous cortisol concentration using the following model:

$$y = \mu+b\beta_{b}+a\beta_{a}+t\beta_{t}+i+\varepsilon$$

where $y$is an $n$ x 1 vector of serum cortisol measurements (μg/dl) at three different timepoints for $n$ samples; $t$ is the timepoint at which the measurement was taken and $\beta_{t}$ is its effect size; $i$ is a random effect term controlling for individual ID; and $b$ and $a$ represent the Elo/behavioral variable and age variable, respectively, consistent with the notations used above.

To analyze diurnal cortisol slope, we used a similar model, but replaced the raw cortisol values in $y$with the slope of serum cortisol from 0800 to 1700 hours μg/dl, added an effect of baseline cortisol measures at 0800, and removed the fixed effect of time point and the random effect (as there were no repeated measures in this analysis).

Finally, for the DEX suppression data, we used three linear models (one for each post-DEX time point) with similar structure as the diurnal cortisol slope model. Here, $y$ was the change in serum cortisol μg/dl from the baseline timepoint (prior to DEX administration) to 1.5, 4.5, or 24 hours post-administration. Because metabolism of dexamethasone can affect the endogenous cortisol response, we controlled for DEX levels at the post-administration timepoint and baseline cortisol levels prior to DEX administration. We also modeled the fixed effects of Elo/behavioral variable (the parameter of interest in this study) and age.

*mtDNA copy number data set*

For mtDNA copy number reported in (7) (data available at https://github.com/ndsimons/MacaqueBehaviorEffects2021), we estimated the effects of Elo and each behavioral variable separately. To do so, we used linear models with the natural log-transformed mtDNA copy number measured for $n$ individuals across up to five flow cytometry-sorted PBMC cell types (helper T cells, cytotoxic T cells, B cells, natural killer cells, and monocytes) as the response variable. In addition to the effects of Elo/behavioral summary and age, we included cell type as a fixed effect predictor variable and a random effect to control for qPCR batch in the mtDNA copy number estimates.

**Table S1.** Data sources and sample sizes.

| **Outcome variable** | **Sample Type** | **N individuals** | **N samples** | **Reference** |
| --- | --- | --- | --- | --- |
| gene expression | flow-sorted helper T cells | 45 | 90 | (1) |
| gene expression | flow-sorted cytotoxic T cells | 45 | 90 |  |
| gene expression | flow-sorted natural killer cells | 45 | 90 |  |
| gene expression | flow-sorted B cells | 44 | 88 |  |
| gene expression | flow-sorted monocytes | 45 | 90 |  |
| gene expression | LPS-challenged white blood cells | 44 | 44 |  |
| gene expression | Baseline white blood cells | 44 | 44 |  |
| gene expression | GARD-challenged white blood cells | 42 | 42 | (5) |
| gene expression | DEX-challenged PBMCs | 43 | 43 | (4) |
| gene expression | DEX-challenged/Baseline PBMCs | 43 | 43 |  |
| chromatin accessibility | DEX-challenged PBMCs | 43 | 43 |  |
| diurnal cortisol concentration | serum | 45 | 45 | (6) |
| diurnal cortisol slope | serum | 45 | 45 |  |
| cortisol conc. 24h post-DEX | 24h post-DEX challenged serum | 45 | 45 |  |
| cortisol conc. 1.5h post-DEX | 1.5h post-DEX challenged serum | 45 | 45 |  |
| cortisol conc. 4.5h post-DEX | 4.5h post-DEX challenged serum | 45 | 45 |  |
| mtDNA copy number | flow-sorted PBMCs | 45 | 45 | (7) |

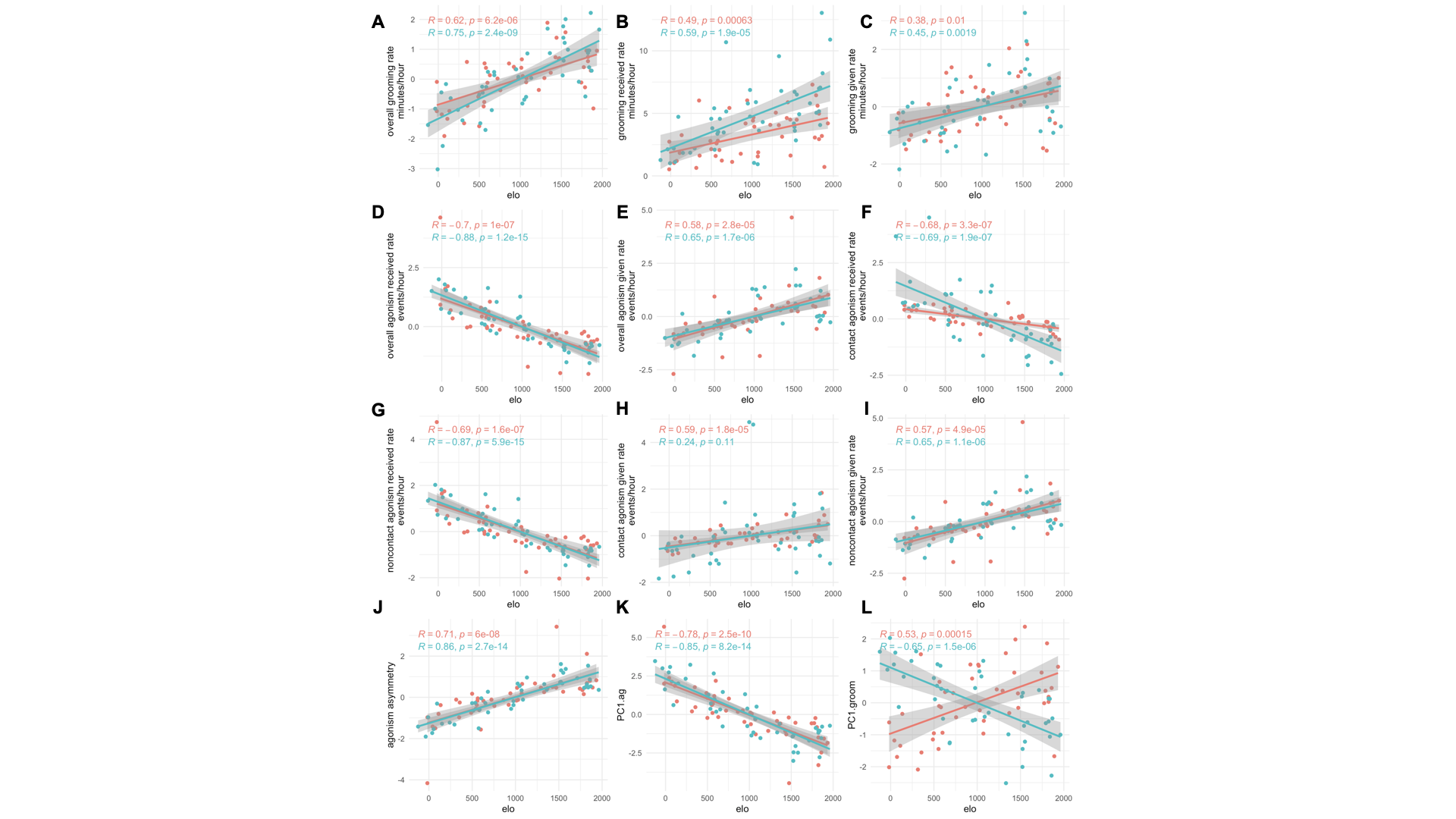

**Figure S1. Relationships between Elo and behavioral variables.** Dominance rank (measured using Elo rating) is correlated with all the behavioral summaries we analyzed here, although the strength of the correlation varies across behavioral variables. In all panels, pink shows the data for Phase 1 and blue shows the data for Phase 2, mean-centered by social group. Note that because PCA imposes arbitrary directionality, PC.1 groom is positively correlated with Elo in Phase 1, but negatively correlated in Phase 2 (Panel L); however, the magnitude of the correlation is similar.

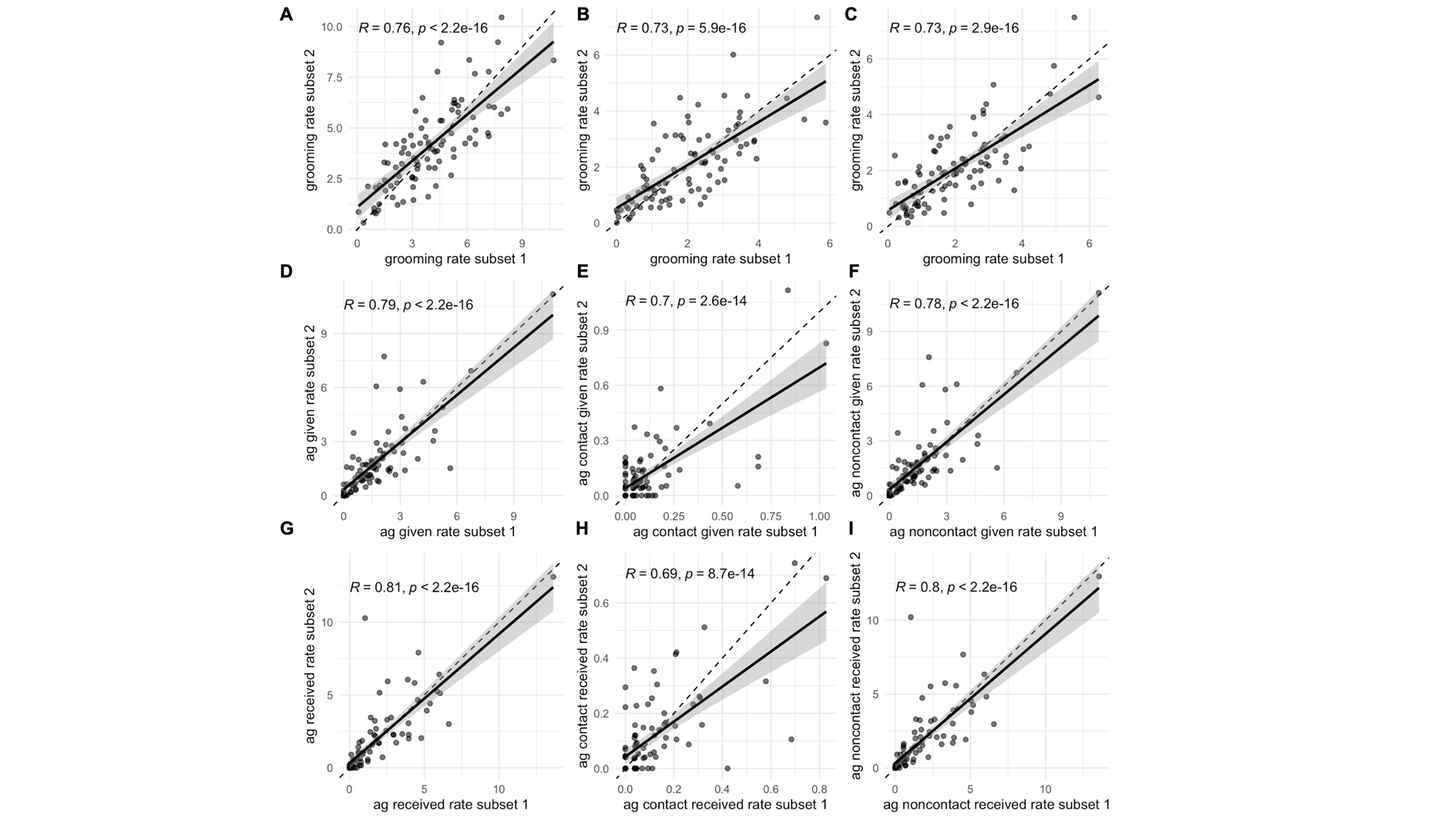

**Figure S2. Behavioral rate estimates are consistent across subsampled data sets**. Each panel shows the correlation between behavioral rates measured in half of the behavioral observation data set against behavioral rates, measured for the same individuals, in the other half of the data sets. Correlations are generally high, suggesting that these rates tend to be consistent across observation blocks, but are lower for measures of agonistic interactions that involve physical contact (E, H), which are relatively rare.

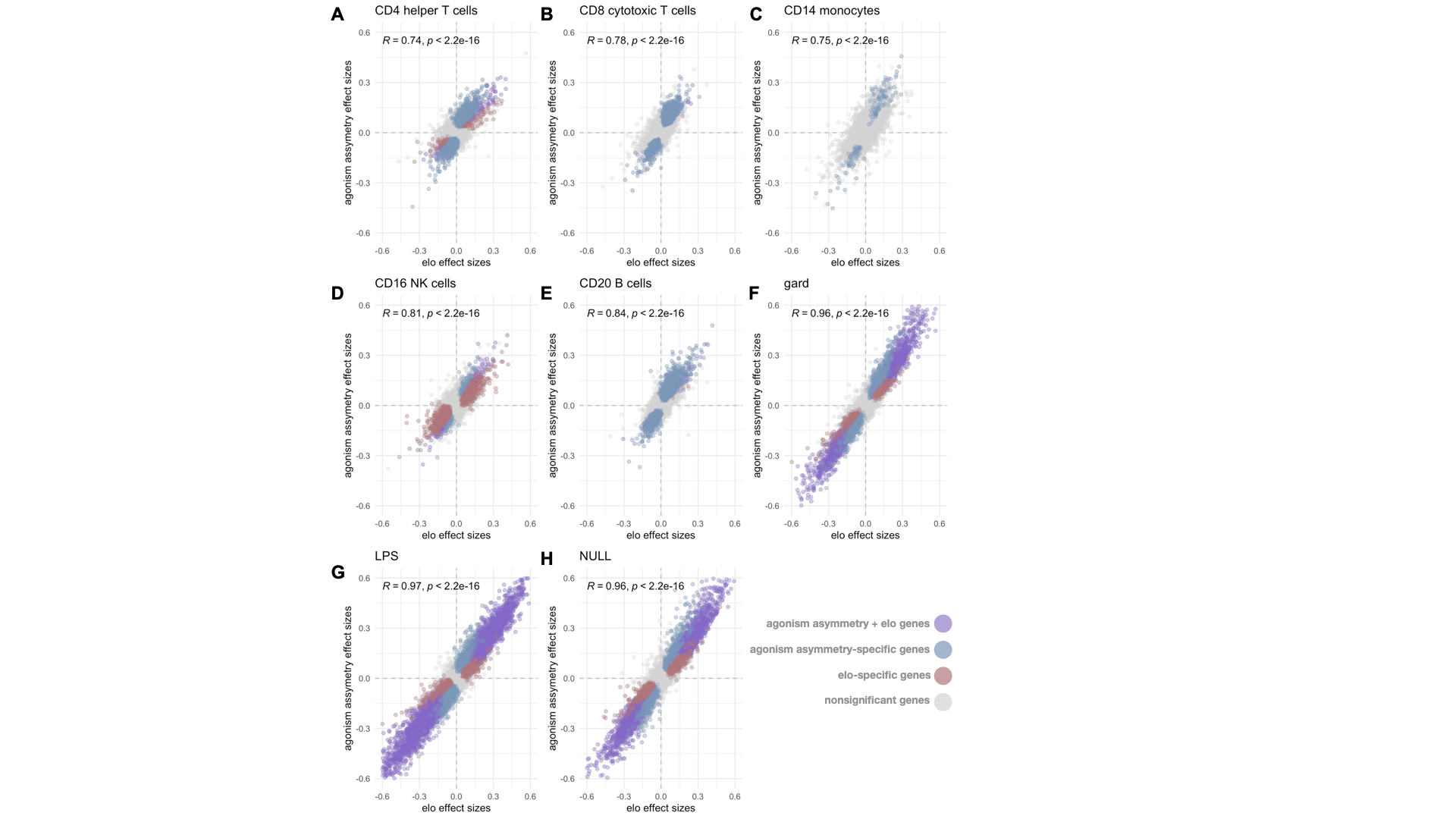

**Figure S3. Effect size correlations between Elo and agonism asymmetry across data sets.** Elo effects and agonism asymmetry effects are highly correlated across all gene expression analyses where agonism asymmetry identified numerous associations, indicating that the increased power to detect agonism asymmetry-associated genes (relative to Elo) reflects a quantitative rather than a qualitative difference in power.

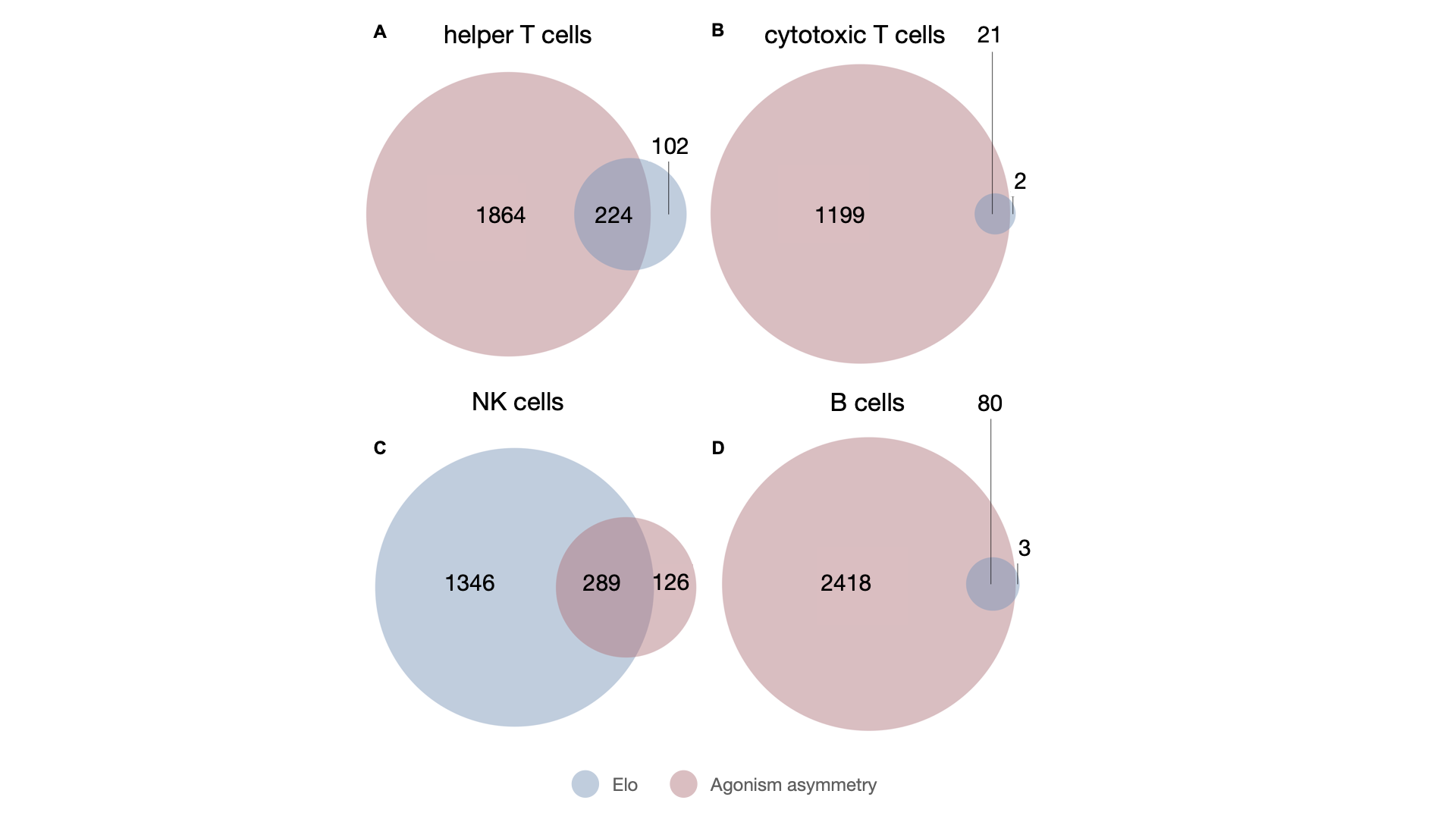

**Figure S4. Overlap between Elo-associated and agonism asymmetry-associated genes.** Each panel shows the overlap between Elo-associated genes and agonism asymmetry-associated genes (all p < 2.2e-16). For three of the four cell types in which we detected Elo effects (all but NK cells), genes associated with agonism asymmetry contain most of the set of genes associated with Elo, but not vice-versa.
